# The dynamics of centromere motion through the metaphase-to-anaphase transition reveal a centromere separation order

**DOI:** 10.1101/582379

**Authors:** Jonathan W. Armond, Katie L. Dale, Nigel J. Burroughs, Andrew D. McAinsh, Elina Vladimirou

## Abstract

During cell division, chromosomes align at the equator of the cell before sister chromatids separate to move to each daughter cell during anaphase. We use high-speed imaging, Bayesian modelling and quantitative analysis to examine the regulation of centromere dynamics through the metaphase-to-anaphase transition. We find that, contrary to the apparent instantaneous separation seen in low-frequency imaging, centromeres separate asynchronously over 1-2 minutes. The timing of separations negatively correlates with the centromere intersister distance during metaphase, which could potentially be explained by variable amounts of cohesion at centromeres. Depletion of condensin I increases this asynchrony. Depletion of condensin II, on the other hand, abolishes centromere metaphase oscillations and impairs centromere speed in anaphase. These results suggest that condensin complexes have broader direct roles in mitotic chromosome dynamics than previously believed and may be crucial for the regulation of chromosome segregation.

## Introduction

One of the crucial tasks of mitosis is to correctly segregate DNA, compactly packaged into chromosomes, to the two daughter cells. Upon mitotic entry, chromatin undergoes a 2-3 fold compaction and a dramatic re-organisation into a higher-order structure to form rod-shaped chromosomes^1–4^. After nuclear envelope disassembly, the chromosomes then attach to the mitotic spindle via kinetochores during prometaphase and congress to the equator of the spindle where they form the metaphase plate. Once all chromosomes are stably attached to the spindle, the spindle assembly checkpoint becomes inactivated and chromatid cohesion is lost as cohesin is cleaved by separase. Finally, sister chromatids separate in anaphase and proceed to their respective spindle poles in a transition that is perceived as a highly synchronous event in human cells^5^.

The condensin protein complexes are fundamental in establishing and regulating chromosome morphology during mitosis^2,6,7^. Eukaryotes have two condensin complexes - condensin I and condensin II - which fulfil non-overlapping roles throughout the cell cycle, are differentially regulated and have differing chromosome binding profiles during the cell cycle^4,8,9^. The two complexes appear all the way along the axis of each chromosome during metaphase, but condensin II is particularly enriched near the inner kinetochore plate^10,11^. Condensins interact with other chromosome scaffold proteins localized along the chromosome axes such as topoisomerase-IΙα (topo-IIα)^12^. Research has predominantly focused on the role of condensins in forming and altering the structure of chromosomes as a precursor of mitosis. How the two condensin complexes influence chromosome dynamics during metaphase, anaphase and the transition between these phases remains largely unknown.

The dynamics of centromere motion during metaphase has been the focus of many quantitative studies^13–19^. The development of automated centromere tracking, advanced image analysis, and data-driven mathematical modelling has provided novel mechanistic and molecular insights into metaphase centromere dynamics ^20,21^. However, the dynamic behaviour of centromeres during the metaphase-to-anaphase transition and during anaphase has been largely overlooked due of the lack of appropriate quantitative tools. Recently the differences in kinetochore phosphorylation states between metaphase and anaphase and how they relate to anaphase chromosome motion has been studied^19^. Here, we have developed a Bayesian statistical model of centromere intersister distance to objectively determine the point of centromere separation via automated tracking of centromeres and to characterise centromere dynamics during and beyond the metaphase-anaphase transition. Using our new assay, we have for the first time quantitatively determined the time distribution of centromere separations in human cells and demonstrated a significant asynchrony: there is a delay of approximately 1-2 minutes between separation of the first and last centromere pairs. This asynchrony is further increased upon condensin I depletion suggesting an important role of the complex in regulating mitotic chromosome dynamics. We show that condensin I has a significant role in timely separation whereas condensin II regulates metaphase chromosome oscillations. Our findings suggest that the two condensin complexes play non-overlapping and critical roles in regulating mitotic chromosome dynamics.

## Results

### Individual centromere separation at anaphase onset is staggered through time

To investigate the dynamic behaviour of chromosomes during the metaphase-to-anaphase transition, we imaged HeLa EGFP-CENP-A cells every 2 s for 5 minutes in 3D (Fig. 1A, S1A), where EGFP-CENP-A labels the centromeres. We also imaged RPE1 GFP-CENP-A / Centrin1-GFP cells, with Centrin1-GFP marking centrosomes (Fig. 1B). We started imaging cells that were already in metaphase and, in many cases, we were able to capture segments of both metaphase and anaphase. Using centromere tracking software^22^, we tracked the position of the centromeres (Figs. 1C-D, S1A) and, in some RPE1 cells, spindle poles (Fig. S1B), through the transition^23^. When we overlaid tracking of the centromere fluorescent markers on movies for visual inspection it was immediately clear that disjunction of sister centromeres was not synchronous at high-temporal resolution. Strikingly, we visually observed a distribution of lag times between the first and last separating centromeres ranging from 1-2 minutes in both HeLa (Fig. 1A) and RPE1 (Fig. 1B) cells. This time delay is significant given that it covers approximately 50% of the duration of anaphase A, the movement of chromosomes toward opposite spindle poles, which is reported to last 3.2±0.8 minutes in HeLa cells^24^.

**Figure 1:**
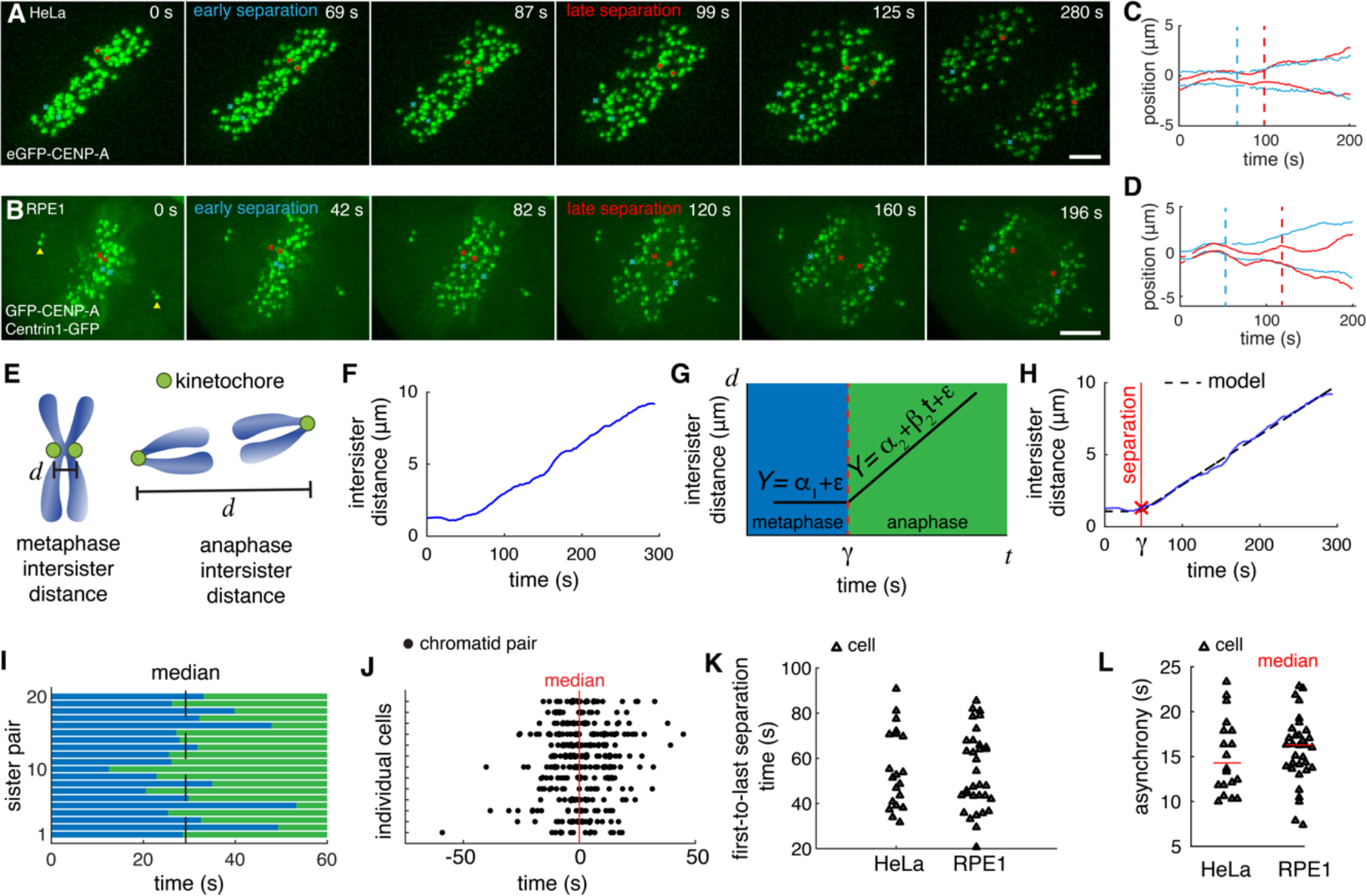
Individual centromere separation at anaphase onset is staggered through time. (A) Images from a movie of HeLa eGFP-CENP-A cell entering anaphase. An early separating (blue) and a late separating centromere pair (red) are shown. Scale bar 3 μm. (B) Images from a movie of an RPE1 GFP-CENP-A Centrin1-GFP cell entering anaphase, with early and late separating centromeres indicated as in A. Centrosomes are shown by yellow triangles. Scale bar 3 μm. (C) Trajectories of the early separating (blue) and late separating (red) centromere pairs marked in A. (D) Trajectories of the early separating (blue) and late separating (red) centromere pairs marked in B. (E) Diagram indicating intersister distance measurement for chromosomes in metaphase and anaphase. (F) Example trajectory of intersister distance during metaphase-to-anaphase transition. (G) Schematic of Bayesian model for metaphase and anaphase components of intersister distance trajectory. Change point between phases is indicated by γ. (H) Example of fitting the Bayesian metaphase to anaphase transition model to the intersister distance trajectory shown in F. Change point is highlighted in red. (I) The transition of individual centromere sister pairs in a single cell from metaphase (blue bars) to anaphase (green bars). (J) Time points of separation of centromeres pairs (circles) in multiple cells (rows). Each cell is aligned by the median separation time at zero. (K) Time difference between first and last separation time for each cell (triangle). (L) Median absolute deviation (MAD) measurement for separation time asynchrony for each cell. n=20 cells for HeLa, n=33 for RPE1.

To obtain an objective quantitative measurement of the centromere separation times, we developed a minimal statistical model of intersister distance dynamics during the metaphase-to-anaphase transition. During metaphase, the distance between sister centromeres (Fig. 1E) grows and shrinks, oscillating around a mean value of 0.88 μm in our HeLa cells, in a process termed chromosome breathing. We measured a coefficient of variation of 29% in intersister distance indicating that the amplitude of the breathing oscillations is relatively small compared to the mean distance (Fig. S1C), hence we can reasonably model metaphase distance using a constant. After separation, centromeres occasionally showed temporary directional reversals (Fig. S1D) as has been previously reported^19,25^, but typically moved consistently towards their respective centrosome (Fig. 1E,F, Fig. S1E).

Therefore, we chose to model the metaphase-to-anaphase transition as a piecewise linear function representing intersister distance as a constant distance during metaphase, followed by a linearly increasing segment during anaphase (Fig. 1G). We employed Bayesian change-point regression^26^ with Markov chain Monte Carlo (MCMC) to fit the model to the trajectory data and locate the change-point γ between metaphase and anaphase. The Bayesian approach allows one to specify prior information, such as approximate feasible ranges for anaphase velocity and metaphase intersister distance, in a principled manner^27^. The model fits well for the majority of intersister distance trajectories (Fig. 1H) during centromere separation and early anaphase, converging with a high *R*^2^ (99.1% of HeLa wild-type trajectories converged during fitting with mean *R*^2^ = 0.85; *n* = 568 converged sister centromere pair trajectories. For RPE1, 99.6% converged with mean *R*^2^ = 0.93 from *n* = 842). Movies of cells with one or more lagging chromosomes, i.e., a chromosome failing to move toward the pole from the metaphase plate for the entire movie, were not included in the analysis. Trajectories that did not converge during the MCMC were discarded (see Methods). This procedure resulted in an average of 15.6 and 17.9 trajectories per cell for HeLa and RPE1, respectively, on par with our previous studies fitting models to automatic centromere tracking^15,28^. Our empirical model enables quantitative study of centromere dynamics during the metaphase-to-anaphase transition by accurately locating the transition timepoint γ itself for individual chromosomes. It could in future be extended to consider the slowing of centromere motion during the later stages of anaphase B, where the two poles start increasing their separation, or be coupled with a more detailed mechanistic model.

Using our metaphase-to-anaphase transition model, we extracted the distribution of centromere separation times per cell. We found that the absolute time of separation of each sister centromere pair varies widely (Fig. 1I, J), with a time delay of 20-90 s between the first and last tracked centromere pair (Fig. 1K). As with all centromere tracking experiments, only a subset of centromeres per cell are successfully tracked, and therefore the time difference between first- and last-detected separating centromere pairs represents a lower bound.

To analyse the distribution of separation times, we calculated a robust estimate for the variation of centromere separation times - median absolute deviation (MAD; see Methods) - to use as a measure of separation asynchrony, with lower values indicating a more synchronous anaphase onset. It is more robust than standard deviation because it is defined around the median rather than the mean and is therefore less sensitive to outliers. Hereafter, we shall refer to MAD as *asynchrony*. The asynchrony plots confirmed that centromere separation times are significantly asynchronous with a median of 14.4 s in HeLa and 16.1 s in RPE1 (Fig. 1L).

To check whether the presence of a lagging chromosome correlates with the degree of asynchrony of the rest chromosomes, we specifically analysed a dataset of cells with at least one lagging chromosome each and, after excluding the centromere of the lagging chromosome from the distribution, we found a range of asynchrony similar to control cells for the rest of the centromeres (Fig. S1F).

### Order and timing of sister centromere separation is not random

An important aspect of centromere asynchronous separation is the order and timing in which successive centromeres separate. It is important to note that this is a distinct notion from the asynchrony measurement which pertains to how the separation events are spread out in time. Having shown that anaphase centromere pair separation is asynchronous, we sought to find if the staggered separation is linked to any spatiotemporal characteristics of the individual centromeres such as their dynamic behaviour or their position within the cell. We therefore tested for correlations *ρ*_*i*_ between the dynamics of centromere sister pairs, their positions within the cell and the timing of their separation (relative to the median) in each cell *i*. We performed a Wilcoxon sign-rank test to determine whether the set of correlations *ρ*_*i*_ was statistically different from zero.

To test whether centromere separation timing correlated with centromere metaphase oscillations, we correlated the magnitude of the autocorrelation at half period, which is an indication of the regularity of oscillations or periodicity, with the separation timing and found a significant positive correlation (Fig. 2A; *p*=0.0051 for HeLa and *p*=0.0039 for RPE1, Wilcoxon sign-rank test), demonstrating that sister centromeres with more regular oscillations separate later. We next examined correlation between average metaphase intersister distance and sister centromere separation timing and, intriguingly, found a negative correlation (Fig. 2B; *p*=0.0013 for HeLa and *p*=0.0016 for RPE1, Wilcoxon sign-rank test). Centromere pairs with a greater inter-centromere distance at metaphase went on to separate earlier than other pairs. We also examined the correlation of separation timing with the centromere’s position within the metaphase plate. The mean distance from the centre of the metaphase plate for each centromere during metaphase significantly correlated with the time of sister centromere separation in both Hela and RPE1 cells (Fig. 2C; *p*=0.019 for HeLa and *p*=1.5×10^−5^ for RPE1, Wilcoxon sign-rank test). This shows that centromeres nearer the periphery of the cell separated earlier. Finally, when we looked at the speed of centromeres in metaphase prior to separating and correlated with separation time, we found no significant correlation with the separation time (Fig. 2D; *p*=0.16 for HeLa and *p*=0.07 for RPE1, Wilcoxon sign-rank test).

**Figure 2:**
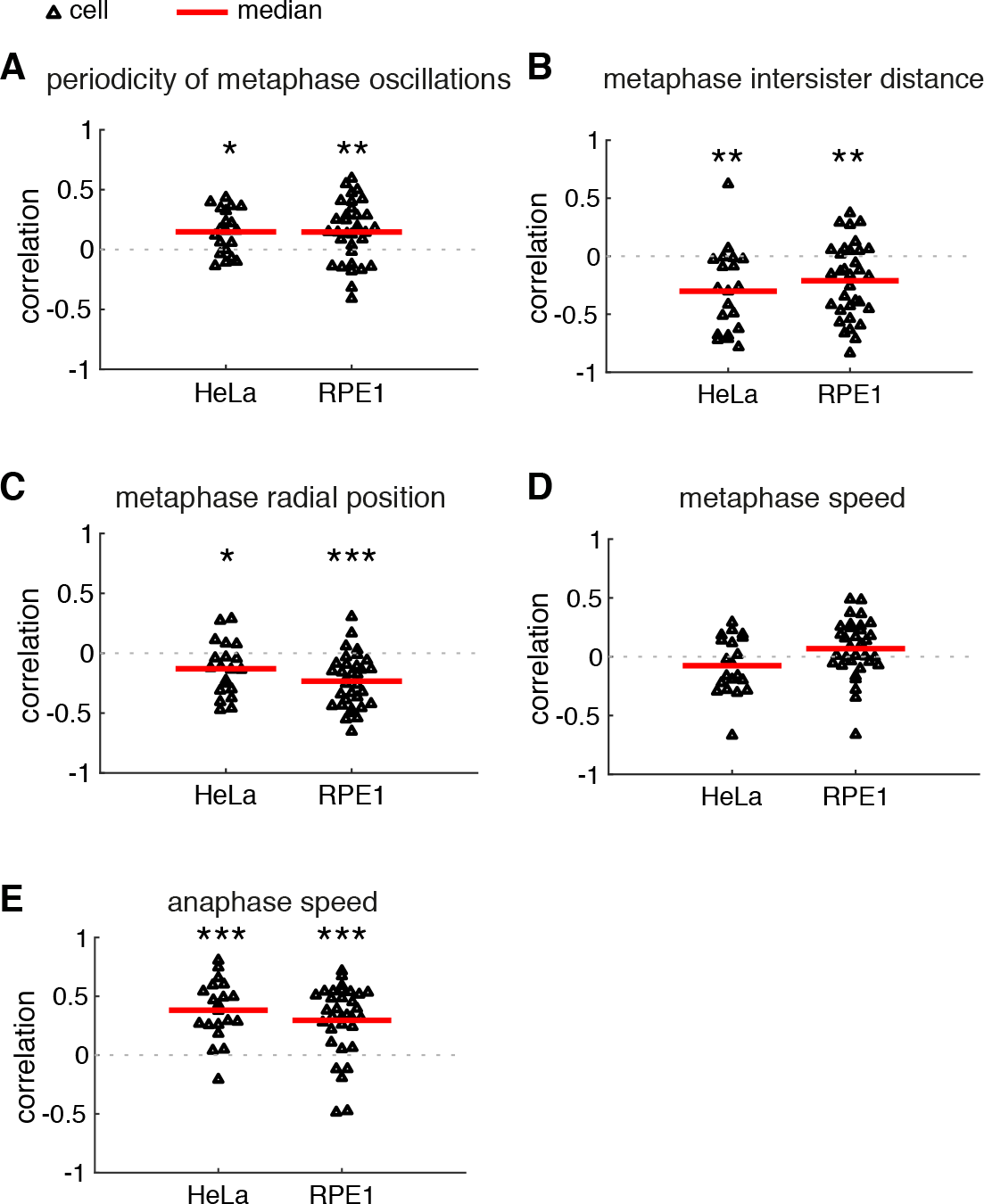
Order and timing of sister centromere separation is not random. Mean correlation per cell (triangle) of centromere pair separation time with (A) periodicity of metaphase oscillations (as per the absolute magnitude of the first negative peak of the autocorrelogram), (B) mean metaphase intersister distance, (C) mean distance during metaphase from the centre of the metaphase plate, (D) mean metaphase speed, (E) mean anaphase speed. Red lines indicate medians across cells. n=20 cells for HeLa, n=33 for RPE1. * is p<0.05, ** is p<0.005, *** is p<0.0005.

Although a *post hoc* event and therefore not a cause of asynchrony, we also tested how centromere speed in anaphase correlates with centromere separation time. There was a significant positive correlation between the speed of centromere motion toward the pole during anaphase and the separation time in both cell types (Fig. 2E; *p*=0.00016 for HeLa and *p*=7.0×10^−5^ for RPE1, Wilcoxon sign-rank test). This demonstrates that the later separating sister centromeres subsequently move toward the pole faster which would lead to an increase in synchrony of arrival at the poles if sustained into anaphase B. However, we cannot determine from our data whether the cause of the increased speed is related to the cause of asynchronous separation or a mechanism to mitigate the asynchrony going into telophase.

### Condensin I is required to maintain minimal centromere separation asynchrony

Having found correlation between metaphase intersister distance and separation timing, we wanted to determine if sister centromere distance was controlling centromere separation asynchrony. We therefore sought to modulate the intersister distance by removing condensins; loss of condensin I significantly increases intersister distances, while loss of condensin II does not^9^. To reduce condensin I and II levels, we depleted the essential subunits CAP-D2 and CAP-D3 respectively by siRNA in HeLa EGFP-CENP-A cells, as previously described^9,29^ (Fig. 3A; referred to as siCondensin-I and siCondensin-II in the following). Under partial depletion of condensins, the cell is able to construct the proper chromosome architecture, as opposed to complete knockout which results in chromosomes collapsing into an amorphous mass^2^. As previously observed^9,30^, the intersister distance was substantially larger when condensin I was depleted compared to control cells (1.3 μm and 0.86 μm for siCondensin-I and siControl, medians respectively, *p*<10^−6^ MW test; Fig. 3B), but depletion of condensin II had a negligible effect (0.88 μm, *p*<10^−6^ MW test; Fig. 3B). Control siRNA treatment resulted in typical centromere trajectories of saw-tooth pattern and frequent directional reversals during metaphase (Figs. 3C and S2A) and siCondensin-I treatment resulted in similar trajectories but with much longer, sustained directed motion before reversing and travelling to the opposite direction (Figs. 3C and S2B), in a manner as previously shown^14,28^. To our knowledge, metaphase centromere motion has not previously been described under condensin II-specific depletion; we observed centromeres that underwent highly irregular directional switching (Figs. 3C and S2C). Using autocorrelation as a measure of centromere oscillatory regularity, we found that siCondensin-II resulted in marked loss of autocorrelation, indicating that centromere motion was no longer periodic (Fig. 3D). Condensin I depletion did not abolish oscillations, but rather caused an increase in the oscillation period (half period 30 s for siControl; 44 s for siCondensin-I; Fig. 3D), confirming our previous work^14,28^. Depletion of condensin II caused a significant reduction in centromere speed in metaphase (1.53 μm min^−1^ and 1.16 μm min^−1^ for siControl, siCondensin-II, medians respectively, *p*<10^−6^ MW test; Fig. 3E). Loss of condensin I caused a statistically significant reduction in centromere speed during metaphase, although the effect was small (1.57 μm min^−1^, *p*<10^−6^ MW test; Fig. 3E). However, both depletions impaired the normal coordination between sisters causing reduced coupling (Fig. 3F).

**Figure 3:**
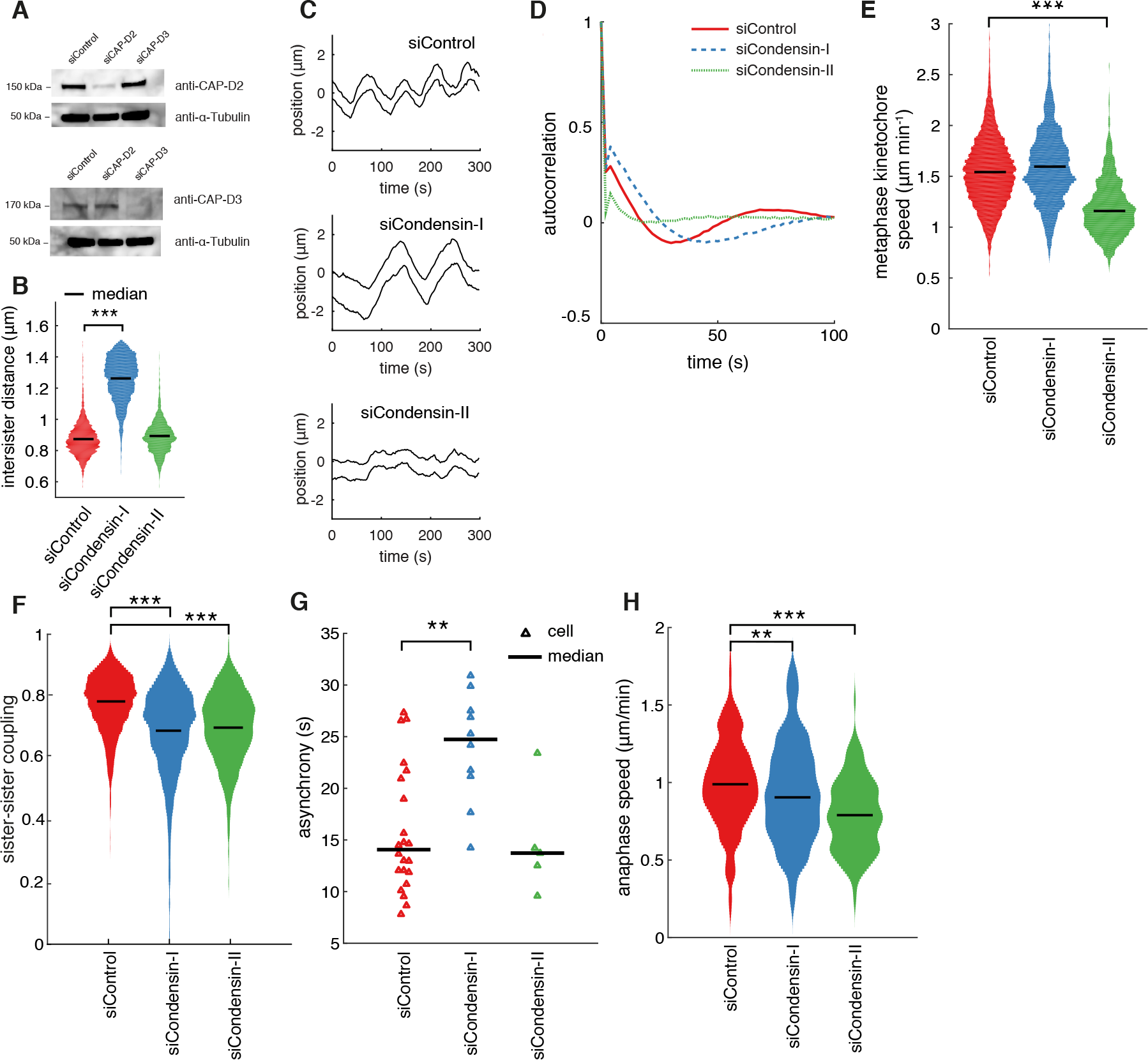
Condensin-II controls centromere metaphase dynamics and condensin-I controls centromere separation asynchrony. (A) Immunoblots of whole cell lysates from HeLa cells transfected with siControl, siCAP-D2 (condensin-I) and siCAP-D3 (condensin-II) oligos and probed with antibodies as indicated 48 hours after transfection. (B) Distribution of mean centromere intersister distances per trajectory. (C) Exemplar trajectories of centromere pair motion in metaphase. (D) Autocorrelation of sister centromere pair centre motion during metaphase. (E) Distribution of mean metaphase centromere speed per trajectory. (F) Distribution of strength of centromere sister-sister coupling as per cross-correlation of sister trajectories during metaphase. (G) Median absolute deviation (MAD) measurement for separation time asynchrony for each cell (triangle). (H) Distribution of mean anaphase speed per trajectory. Black lines indicate medians. * is p<0.05, ** is p<0.005, *** is p<0.0005. n=8371 centromere pairs for siControl, n=2183 for siCondensin-I, n=2076 for siCondensin-II. In G, n=22 cells for siControl, n=10 for siCondensin-I and n=5 for siCondensin-II.

We then calculated asynchrony and found depletion of condensin I had the striking effect of increasing centromere separation asynchrony by over 50% (Fig. 3G; *p*=0.002 MW test). Condensin II depletion, with unchanged intersister distance, did not affect this asynchrony (Fig. 3G). When we looked at the speed of centromeres in anaphase, we found a small difference between siControl and siCondensin-I (0.99 μm min^−1^ and 0.91 μm min^−1^ for siControl, siCondensin-I, medians respectively, *p*=0.004 MW test; Fig. 3H), but a much larger drop under siCondensin-II (0.78 μm min^−1^, *p*<10^−6^ MW test; Fig. 3H). Taken together, our findings show that the two condensin complexes have non-overlapping roles in regulating mitotic chromosome dynamics. While condensin II plays a previously unreported role in setting centromere metaphase dynamics, condensin I has a significant effect on centromere separation dynamics at the metaphase-to-anaphase transition.

Having found that depletion of condensin I caused a dramatic increase in asynchrony, we sought to test if the negative correlation between the metaphase intersister distance and separation timing we found in unperturbed cells held true. Surprisingly, we found that when condensin I was depleted the correlation became inverted so that larger intersister distances correlated with later separation instead of earlier (*p*=0.0014 MW test; Fig. 4A). All other correlations were not significantly different from control cells (Fig. 4B-E).

**Figure 4:**
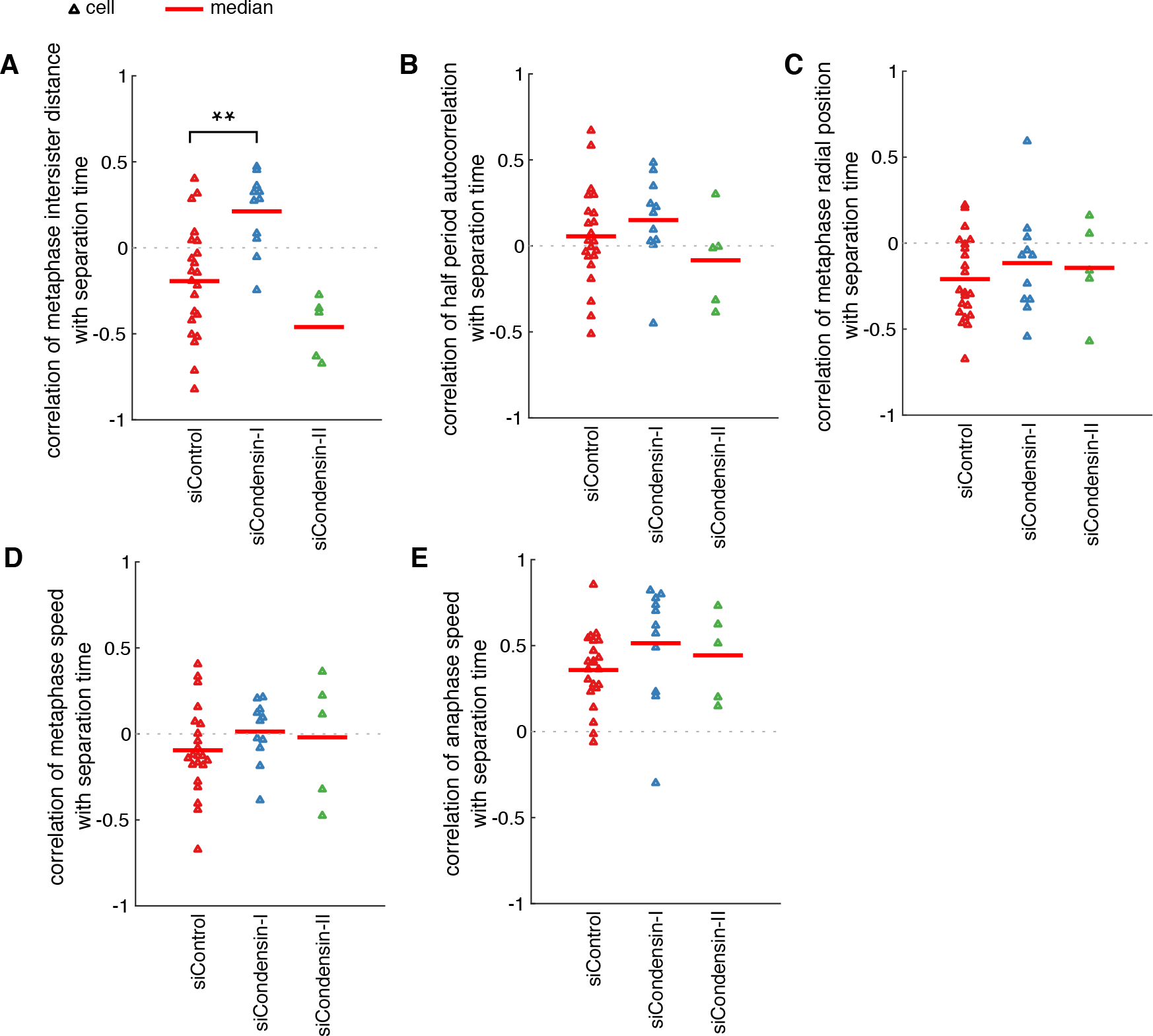
Order and timing of sister centromere separation under condensin depletions. Mean correlations per cell under depletions of centromere pair separation time with (A) mean metaphase intersister distance, (B) periodicity of metaphase oscillations (as per the absolute magnitude of the first negative peak of the autocorrelogram), (C) mean distance during metaphase from the centre of the metaphase plate, (D) mean metaphase speed, (E) mean anaphase speed. Red lines indicate medians across cells. * is p<0.05, ** is p<0.005, *** is p<0.0005. n=22 cells for siControl, n=10 for siCondensin-I and n=5 for siCondensin-II.

One of the functions of condensin I is to target topo-IIα^12^ which is responsible for the resolution of catenation of sister chromatid DNA^12,31–34^. Catenation is a consequence of DNA replication^35^. While the majority of catenanes found along the chromosome arms are resolved before metaphase by the activity of topo-IIα^12,31–35^, centromere catenation has been shown to persist into anaphase^36^. To determine whether the condensin I phenotypes we observed were due to topo-IIα mislocalisation, we treated cells with ICRF-187, a highly specific inhibitor of topo-IIα^37,38^. We imaged cells 1-3 hours post-treatment as in previous studies^36,38^; after 3 hours, most mitotic cells were arrested in metaphase. The treated cells did not recapitulate the phenotypes associated with condensin I depletion, or indeed condensin II, instead showed normal metaphase intersister distance, speed and oscillations (Fig. S3A-E). Moreover, ICRF-187 treatment presented a normal degree of asynchrony (Fig. S3F) and poleward velocity (Fig. S3G). This demonstrates that the observed disruption of centromere separation timing upon condensin I depletion is not due to downstream effects on topo-IIα.

## Discussion

It is well known that a single unattached kinetochore, by virtue of binding Mad2, can prevent anaphase via the spindle assembly checkpoint (SAC)^5^ – our data show that each centromere pair can take a different amount of time to start moving poleward after the first separation. Here, we provide the most comprehensive quantitative analysis to date of centromere separation asynchrony at the metaphase-anaphase transition in living human cells. This asynchrony spans over 1-2 minutes and is linked to intersister centromeric distance.

Early observations using fixed samples of metaphase spreads of untreated or colcemid-arrested cells from plants, Chinese hamster, Ptk1 and human leukocytes stained with Giemsa, suggested that some chromosomes separate earlier than others in the centromeric region and that there might be a pattern of separation with chromosomes 1 and 2 separating earlier^39,40^. However, these observations could not provide quantitative timing data having been performed on fixed cells. To our knowledge, there are two studies which observed centromere separation asynchrony in live cells but were limited either by temporal resolution or by following only two chromosomes. Time-lapse DIC microscopy observations of *Haemanthus katharinae* cells revealed that some chromosomes may start anaphase movement up to 1 minute later than others, although usually the difference is less than 30 seconds^41^. In *S. cerevisiae*, the median time between the two chromosome segregation events was found to be around 90 s and also chromosome 4 always segregated before chromosome 5^42^. Taken together with our results, centromere separation asynchrony and a degree of ordering appears to be a conserved phenomenon occurring across many species during cell division.

Our data show that the centromere asynchrony is significantly increased when cells are depleted of condensin I, but not of condensin II. Previous studies have shown that depleting condensin I or condensin II results in similar degrees of chromosome bridges and irregular chromosomes^9^ and therefore the observed increased asynchrony upon condensin I depletion, but not condensin II, could not be explained by the presence of chromosome bridges. Neither could the presence of DNA ultrafine bridges (UFB) explain the observed increased asynchrony upon condensin I depletion because inhibition of topo-IIα, which is responsible for resolving DNA catenations that underlie UFB formation, did not result in an increase in sister centromere separation asynchrony.

Our results raise an interesting question: how does centromere intersister distance relate to the timing of separation? Although in principle there can be numerous reasons why some chromosomes have higher intersister distance, it seems most likely to be caused by lower centromeric cohesion. Cohesion is maintained by both cohesin rings and catenation^36^. The sizes of human centromeres vary^43^ and it seems plausible that smaller centromeres would bind fewer cohesin rings or harbour less catenation. Having less of either, resulting in reduced cohesion, could conceivably cause faster sister separation (Fig. 5).

**Figure 5.**
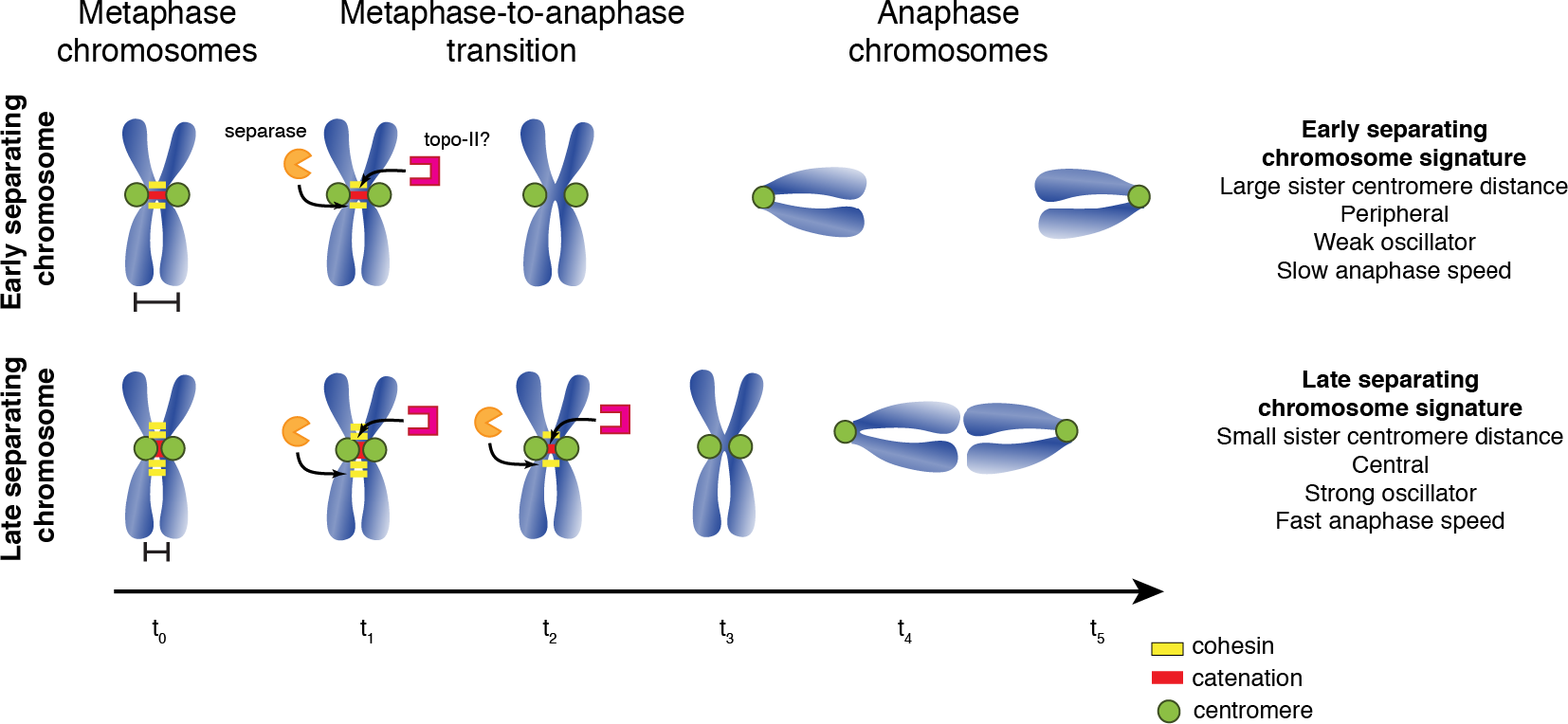
Working model for centromere separation asynchrony. A possible model of how centromere intersister distance controls centromere separation timing. Chromosomes with larger intersister distance have less cohesion – mediated by cohesin and catenation – which is removed faster and hence separate earlier (top). Those with shorter distance take comparatively longer to initiate separation (bottom). Early separating chromosomes are found in the periphery of the cell, are weak metaphase oscillators, and exhibit slow anaphase speed. Late separating chromosomes are found more centrally within the metaphase plate, are strong metaphase oscillators, and exhibit fast anaphase speed.

Condensin I was shown to be required for the complete dissociation of cohesin from chromosome arms^29^. Our observed increased asynchrony upon condensin I depletion could suggest that condensin I is also required for mediating the removal of centromeric cohesin at the metaphase-to-anaphase transition. Moreover, removal of cohesin may be a prerequisite for centromere decatenation^35^. Alternatively, condensin I might be directly involved in centromere decatenation as suggested by experiments in budding yeast, where complete sister chromatid decatenation of circular minichromosomes required condensin activity, in a topo-IIα independent manner^44^.

Our analysis demonstrates a negative correlation between the timing of sister centromere separation and metaphase centromere intersister distance in untreated cells. However, depletion of condensin I reversed the correlation between the timing of sister centromere separation and the metaphase intersister distance; the two measurements were positively correlated as pairs with larger intersister distance separated later. A speculative explanation for this paradoxical result, is that above a certain threshold intersister distance becomes detrimental to a quick separation because the tension across the centromere sister pair at such distance is so high that the kinetochore-microtubule fibres become stabilised. Indeed, pulling force on kinetochore-microtubule fibres is known to reduce likelihood of depolymerisation^45,46^. Evidence pointing in this direction is given by metaphase centromeres in HeLa cells, where small but statistically significant negative correlations between both intersister distance and metaphase centromere speed (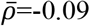, *p*=1.6×10^−6^), and intersister distance and periodicity (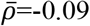, *p*=1.2×10^−6^), indicate less kinetochore-microtubule fibre activity on centromere pairs with higher intersister distance.

Although condensin II depletion left asynchrony unchanged, we did find that oscillation regularity was severely compromised while metaphase and anaphase centromere speeds were reduced. While previous studies have shown that depletion of condensin II does not affect mitotic progression^29^, other studies have shown that the kinetochore-microtubule interactions are compromised in condensin-depleted chromosomes, with the back-to-back orientation of sister kinetochores being destabilised^10^. Indeed, depletion of both condensin complexes via SMC2 depletion, the core subunit of both condensin complexes^8^, resulted in elongated kinetochores which are able to attach to microtubules but are defective in forming stable amphitelic attachments^2^. Taking together previous reports of the distinctive localisation of condensin II at the centromere near the inner kinetochore and our metaphase and anaphase centromere dynamics results, we hypothesise that it is the depletion of condensin II that results in defective kinetochore structure, altering kinetochore-microtubule dynamics. Moreover, the lack of effect on asynchrony suggests that kinetochore-microtubule interactions are unlikely to be inducing separation timing differences.

In summary, we show that the two condensin complexes make distinct contributions to the establishment of normal metaphase and anaphase centromere dynamics, in addition to the well-established roles in maintaining and modifying chromosome structure^29^. In particular, our results point to a previously unknown role for condensin II complexes in metaphase centromere dynamics that is critical for normal oscillatory motion. However, only condensin I is required for timely centromere separation. What benefit could a cell derive from regulating the synchrony and order of centromere separation? One hypothesis is that it may help to ensure that chromosomes reach the appropriate position with the correct temporal order to help maintain a spatial order that is transmitted across the cell cycle^47^. Deciphering chromosome-specific dynamics and chromosome spatiotemporal organisation throughout the cell cycle by integrating quantitative live cell imaging, uniquely labelled chromosomes and data-driven modelling is an important area for further study.

## Methods

### Cell culture, small interfering RNA transfection and drug treatments

HeLa K EGFP–CENP-A were cultured in DMEM-GlutaMAX medium containing 10% FBS, 100 units/mL penicillin and 100 µg/mL streptomycin and 0.5 μg/ml Puromycin. RPE1 GFP-CENP-A / Centrin1-GFP cells (kind gift from A. Khodjakov) were cultured in DMEM/F-12 medium containing 2mM L-Glutamine, 10% FBS and 100 units/mL penicillin and 100 µg/mL streptomycin. All cell cultures were maintained at 37ºC with 5% CO2 in a humidified incubator. Small interfering RNA (siRNA) oligonucleotides (Invitrogen) CAP-D2 (5’-CCAUAUGCUCAGUGCUACATT-3’), CAP-D3 (5-CAUGGAUCUAUGGAGAGUATT-3)^29^ and control^15^ were transfected using Oligofectamine transfection reagent (Invitrogen) according to the manufacturer’s instructions and analysed 48h after transfection. For drug treatments, cells were treated with the Topoisomerase-II inhibitor Dexrazoxane (ICRF-187, Sigma-Aldrich) for 1 hour at 5 μM final concentration and DMSO (Sigma-Aldrich) prior to imaging.

### Immunoblotting

Cells were collected by centrifugation, cell pellets were washed in PBS, and lysed in mammalian protein extraction reagent M-PER (Thermofisher) supplemented with Halt protease and phosphatase inhibitor cocktail (Thermofisher). Protein concentration was determined using the Direct Detect assay (Millipore). Protein samples were run on 4–15% Mini-PROTEAN TGX gels (Bio-Rad), and transferred onto nitrocellulose membranes (Thermofisher). Membranes were incubated with the following primary antibodies: anti-CNAP1 (NCAPD2) Rabbit mAb (Abcam, ab137075); anti-NCAPD3 Rabbit mAb (Cell Signalling Technology, 13473); anti-α-Tubulin Mouse mAb (Thermofisher, 32-2500); and horseradish peroxidase-conjugated secondary antibodies (1:10000, GE Healthcare). Protein bands were visualised using SuperSignal™ West Pico PLUS Chemiluminescent Substrate (Thermofisher), and an ImageQuant LAS 4000 image analyser (GE Healthcare).

### Live cell imaging and centromere tracking

Cells were imaged in Leibovitz L-15 (Gibco) supplemented with 10% FBS and imaged using a 100X 1.4NA oil objective on a Perkin-Elmer VOX Ultraview spinning disk confocal microscope with a Hamamatsu ORCA-R2 camera. For metaphase analysis, we captured 25 Z-planes over 12.5 μm Z-range every 2 seconds for 5 mins. All movies were visually inspected and cells exhibiting gross errors or impairments were discarded. In particular, all metaphase-to-anaphase transition movies showing a lagging chromosome, i.e., a chromosome failing to move toward the pole from the metaphase plate throughout the whole movie, were not included in the analysis. Occasionally, a non-disjunction of a centromere pair was observed. In these cases, we discarded the trajectory of that pair, but did not discard the whole cell. Centromeres were tracked using the KiT^22^ software v1.5 in MATLAB, but we did not use the metaphase plate to define a coordinate system as previously, instead measuring the Euclidean distance between centromere pairs. Assignment of poles was by identifying the extreme outlying fluorescent spots in appropriate positions. Centromere trajectories from cells exclusively in metaphase were analysed using the CupL software in MATLAB^28^.

### Statistical methodology

We computed all tests in MATLAB and report p-values as either *p*<10^−6^ or as the exact value if p>10^−6^. The type of test used is noted in the text. We used a robust statistic analogous to standard deviation but less sensitive to outliers – median absolute deviation (MAD) – to describe the asynchrony of centromere separation times. This is defined as mad(X) = median(|*X*_*i*_ − median(*X*)|).

### Centromere separation model

We sought to establish a model-based quantitative analysis of anaphase centromere separation dynamics. However, in anaphase it was not possible to use the metaphase plate as a reference frame to compensate for translation and rotation of the cell because the plate separates at anaphase. The spindle poles were also not reliable reference points due to following a somewhat independent motion with respect to the centromeres. This was confirmed by the low correlation between rotations of the pole-to-pole axis and metaphase plate normal axis (*ρ*=0.11) and suggests flexibility within the spindle.

Therefore, we performed all analyses on Euclidean distance measurements between pairs of centromeres, which have the advantage of being invariant under rotation or translation. We modelled the distance *Y*_*i*_ between sister centromeres at time *t*_*i*_ during the metaphase-to-anaphase transition using a two segment piecewise linear Bayesian regression: *Y*_*i*_ = *α*_1_ + *ε*_*i*_ for *t*_*i*_ < *γ* and *Y*_*i*_ = *α*_2_ + *β*_2_*t*_*i*_ + *ε*_*i*_ for *t*_*i*_ ≥ *γ*, where *γ* is the change point between segments and continuity is imposed by the condition *α*_1_ = *α*_2_ + *β*_2_*γ*. Here, *γ* is the crucial parameter representing the timepoint of centromere separation. We used a non-informative uniform prior on *γ* and weakly informative priors on all other parameters (see Supplementary Text 1 for full details). We developed a Metropolis-within-Gibbs sampler in MATLAB to generate posterior samples, running for 2000 iterations and discarding the first 500 iterations. Convergence was checked visually and by calculating the Gelman-Rubin potential scale reduction statistic over four independent chains^48^.

## Acknowledgements

The authors thank Dr. Sarah McClelland (Barts Cancer Institute, London, U.K.) and Dr. Rob Hynds (UCL Cancer Institute, London, U.K.) for helpful discussions and critical reading of the manuscript and Prof. Alexey Khodjakov (Wadsworth Center, Albany, N.Y., U.S.) for providing the RPE1 GFP-CENP-A, centrin1-GFP cell line. E.V. is supported by a Cancer Research UK Lung Cancer Centre of Excellence Supplementary Award (C7893/A24956). A.D.M. is supported by a Wellcome Trust Senior Investigator Award (106151/Z/14/Z) and a Royal Society Wolfson Research Merit Award (WM150020).

## Author contributions

E.V. and A.D.M. planned and supervised research. J.W.A. and E.V. performed all experiments apart from the western blots which were performed by K.L.D.. J.W.A. and E.V. developed the chromosome separation model and performed all the data analysis. J.W.A., and E.V. wrote the paper with contributions from A.D.M., N.J.B and K.L.D..

