## Supplemental Material for "The dynamics of centromere motion through the metaphase-to-anaphase transition reveal a centromere separation order"

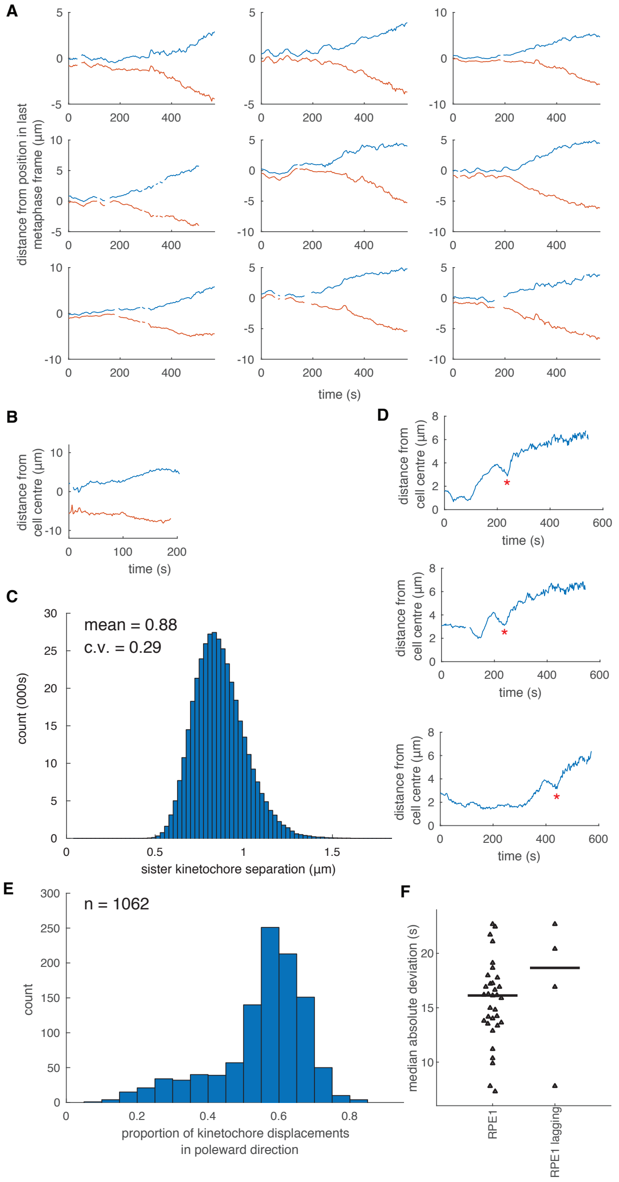


**Figure S1. Tracking centromeres through the metaphase-to-anaphase transition.** (A) Example trajectories of separating centromere pairs transitioning from metaphase through to anaphase. (B) Example trajectory of pole (centrosomes) separation during metaphase-to-anaphase transition. (C) Histogram of all measurements of sister-sister centromere distance in metaphase. (D) Example centromere separation trajectories exhibiting temporary direction reversals. (E) Proportion of frames from each centromere trajectory where the centromere moved in a poleward direction. (F) MAD of RPE-1 cells (triangles) with no lagging chromosomes and those with lagging chromosomes.

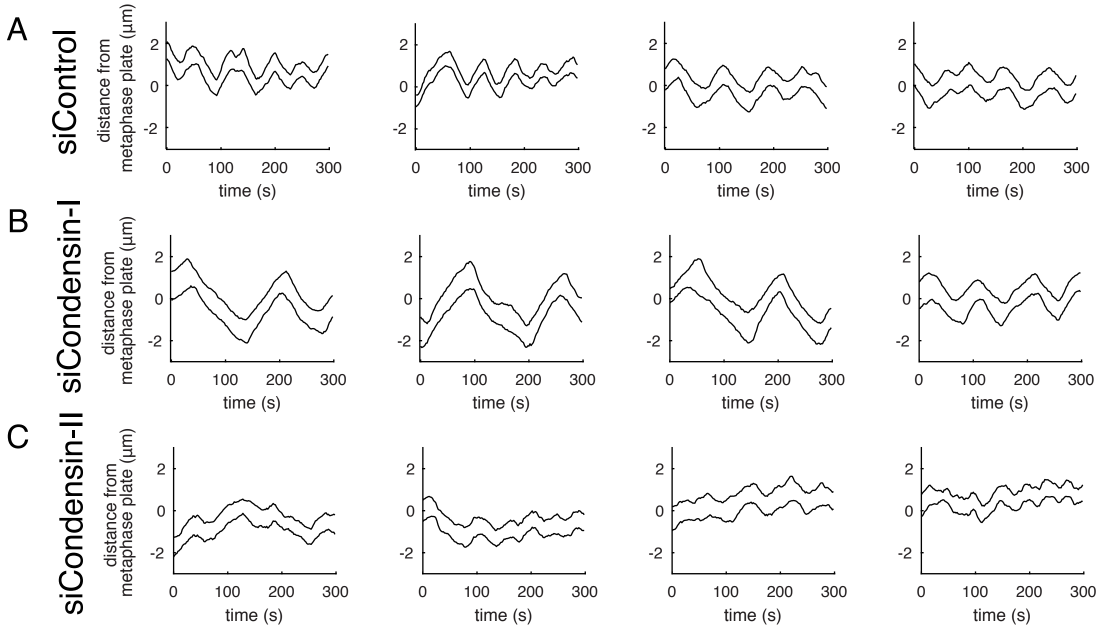


**Figure S2. Representative examples of sister centromere pair trajectories.**
Example centromere trajectories from HeLa cells transfected with (A) siControl, (B) siCondensin-I, (C) siCondensin-II.

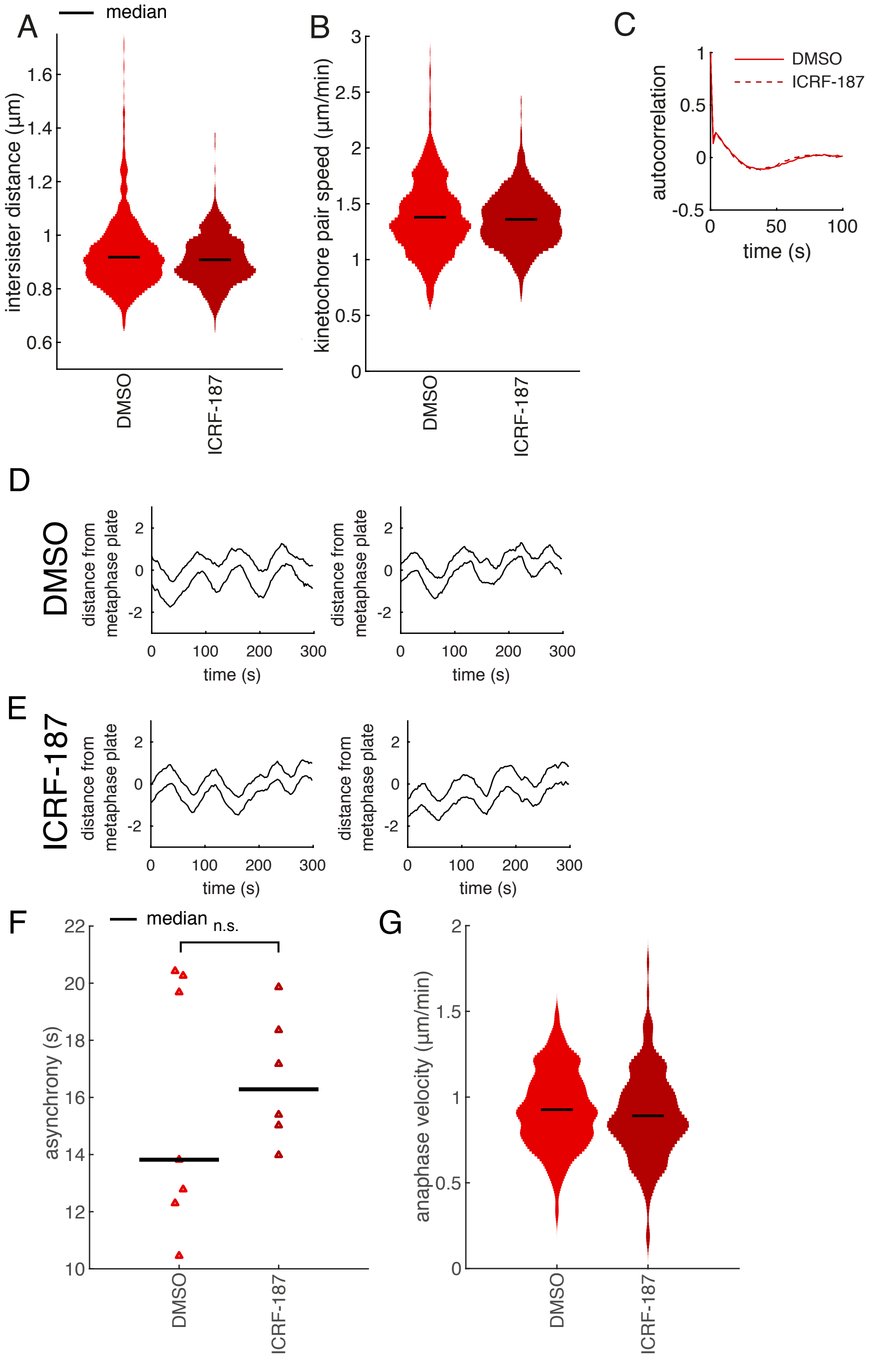


**Figure S3. Centromere dynamics of cells under topoisomerase-IIα inhibition.**

Cells treated with ICRF-187 (5 µM final concentration), a topo-IIα inhibitor or DMSO control. (A) Distributions of mean centromere intersister distance of HeLa cells per trajectory. (B) Distributions of mean centromere pair speeds per trajectory. (C) Average autocorrelation of centromere pair displacements. Representative examples of sister centromere pair trajectories from HeLa cells treated with (D) DMSO and (E) ICRF-187. (F) MAD for cells (triangles) treated with DMSO and ICRF-187, black lines indicate medians. (G) Distribution of anaphase velocity. Black lines indicate medians. n=1175 trajectories for DMSO and n=714 for ICRF-187. In F, n=7 cells for DMSO and n=6 for ICRF-187.

### Supplementary Text 1

#### Bayesian detection of centromere separation

Let $Y_i$ be the inter-sister distance at time point $t_i$. We model $Y_i$ as a piecewise function of one constant (metaphase) and one linear part (anaphase), with the change point at $t = \gamma$, as follows

$$Y_i = \alpha_1 + \epsilon_i \qquad t_i \leq \gamma \tag{1}$$
$$Y_i = \alpha_2 + \beta_2 t_i + \epsilon_i \quad t_i > \gamma \tag{2}$$

where $\epsilon_i$ are i.i.d. normal errors with variance $\sigma^2$. Note $\gamma$ is a continuous parameter and not necessarily equal to any of $t_i$. We impose continuity between (1) and (2), by fixing $\alpha_1 = \alpha_2 + \beta_2\gamma$. Hence the data are distributed normally

$$Y_i \sim \mathcal{N}(\alpha_1, \sigma^2) \qquad i \leq k \tag{3}$$
$$Y_i \sim \mathcal{N}(\alpha_2 + \beta_2 t_i, \sigma^2) \qquad i > k \tag{4}$$

Following [3] we used Bayesian methods to infer location of the change point $\gamma$ and parameters $\alpha_2, \beta_2, \sigma^2$. The Bayesian methodology has the advantage of being conceptually simple in implementation and interpretation, and allows the incorporation of prior knowledge [1]. Thus we seek the posterior distribution

$$p(\gamma, \alpha_2, \beta_2, \tau|\mathbf{y}) \propto p(\mathbf{y}|\gamma, \alpha_2, \beta_2, \tau)p(\gamma)p(\alpha_2)p(\beta_2)p(\tau) \tag{5}$$

where $\mathbf{y} = (y_1, \ldots, y_n)$ is a realization of $\mathbf{Y} = (Y_1, \ldots, Y_n)$ and $\tau = 1/\sigma^2$ parameterizes $\sigma^2$. The likelihood of the data under this model is given by

$$\mathcal{L}(\gamma, \alpha_2, \beta_2, \tau) = p(\mathbf{y}|\gamma, \alpha_2, \beta_2, \tau) = \frac{\tau^{n/2}}{(2\pi)^{n/2}} \exp\left\{-\frac{\tau}{2}\left[\sum_{t\leq\gamma}(y_i - \alpha_1)^2 + \sum_{t>\gamma}(y_i - \alpha_2 - \beta_2 t_i)^2\right]\right\} \tag{6}$$

We imposed a noninformative prior on $\gamma$ since we have no prior information on where the change will occur

$$\gamma \sim \text{Uniform}(0, t_n) \tag{7}$$

and weakly informative priors, based on physiologically reasonable limits, on the remaining parameters

$$\tau \sim \text{Gamma}(c, d) \tag{8}$$
$$\alpha_2 \sim \text{Normal}(\mu_\alpha, \sigma_\alpha^2) \tag{9}$$
$$\beta_2 \sim \text{Folded-Normal}(\mu_\beta, \sigma_\beta^2) \tag{10}$$

where the hyperparameters are $c = 2$, $d = 5$, $\mu_\alpha = 0$, $\mu_\beta = 1/60$, and $\sigma_\alpha = \sigma_\beta = 100$. We used a folded-normal for $\beta_2$ since intersister distance always tends to increase after centromere separation.

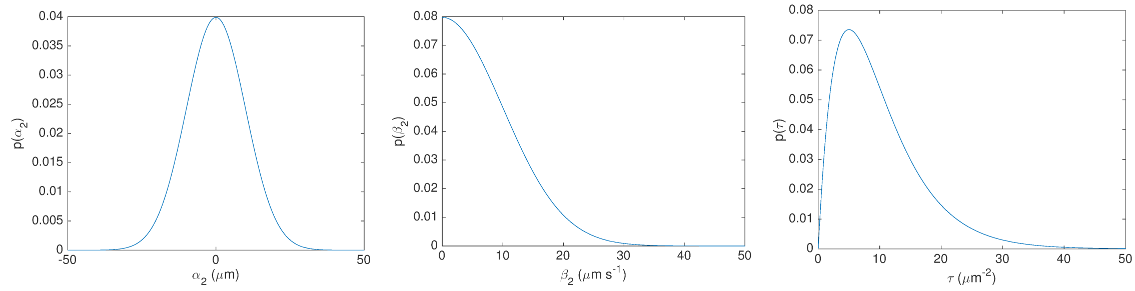


(a) Prior for $\alpha_2$ (b) Prior for $\beta_2$ (c) Prior for $\tau$

Following [4], we implemented a Gibbs sampler to generate posterior samples with the following full conditionals

$$p(\tau|\cdot) \propto \text{Gamma}\left(\frac{n}{2} + c, d + \frac{1}{2}\sum_{t\leq\gamma}(y_i - \alpha_1)^2 + \frac{1}{2}\sum_{t>\gamma}(y_i - \alpha_2 - \beta_2 t_i)^2\right) \tag{11}$$

$$p(\alpha_2|\cdot) \propto \text{Normal}\left(\tau_{\alpha_2}^{-1}\left(\mu_\alpha + \tau\sigma_\alpha^2\sum_{t>\gamma}(y_i - \beta_2 t_i)\right), \frac{\sigma_\alpha^2}{2\tau_{\alpha_2}}\right) \tag{12}$$

$$p(\beta_2|\cdot) \propto \text{Folded-Normal}\left(\tau_{\beta_2}^{-1}\left(\mu_\beta - \tau\sigma_\beta^2\sum_{t>\gamma}t_i(\alpha_2 - y_i)\right), \frac{\sigma_\beta^2}{\tau_{\beta_2}}\right) \tag{13}$$

where $\tau_{\alpha_2} = (n-k)\tau\sigma_\alpha^2 + 1$ and $\tau_{\beta_2} = 1 + \tau\sigma_\beta^2\sum_{t>\gamma}t_i^2$, with $k = \sup\{i : t_i \leq \gamma\}$. Non-negativity was imposed on $\beta_2$ by rejection sampling. We sampled $\gamma$ by rejection sampling with proposals drawn by inverse discrete transform sampling from a piecewise uniform density constructed by maximizing conditional posterior $p(\gamma|\cdot)$ between each datum, as suggested in [4]. This amounts to setting $\gamma = (\sum_{i=1}^n Y_i - j\alpha_2)/(j\beta_2)$ in the likelihood $\mathcal{L}$ for each between-datum segment $t \in [t_j, t_{j+1})$ and normalizing the resulting distribution.

We ran the sampler for 2000 iterations in the first instance, discarding the first half as burn-in and thinning every 2 samples, resulting in 500 samples. We checked convergence computing the Gelman-Rubin potential scale reduction $\sqrt{\hat{R}}$ statistic [2] on 5 chains with a convergence criteria of $\sqrt{\hat{R}} < 1.1$. For trajectories that did not converge we ran the sampler again, doubling the number of iterations and thinning interval. We repeated this extension process a further third time, if necessary, before discarding the trajectory.
